# Associative learning in larval and adult Drosophila is impaired by the dopamine-synthesis inhibitor 3-Iodo-L-tyrosine

**DOI:** 10.1101/2020.10.26.354688

**Authors:** Juliane Thoener, Christian König, Aliće Weiglein, Naoko Toshima, Nino Mancini, Fatima Amin, Michael Schleyer

## Abstract

Across the animal kingdom, dopamine plays a crucial role in conferring reinforcement signals that teach animals about the causal structure of the world. In the fruit fly *Drosophila melanogaster*, the dopamine system has largely been studied using a rich genetic toolbox. Here, we suggest a complementary pharmacological approach applying the dopamine-synthesis inhibitor 3-Iodo-L-tyrosine (3IY), which causes acute systemic inhibition of dopamine signaling. Using Pavlovian conditioning, across developmental stages (3^rd^ instar larva versus adult), valence domains (reward versus punishment), and types of reinforcement (natural versus optogenetically induced), we find that 3IY feeding specifically impairs associative learning, whereas additional feeding of L-3,4-dihydroxyphenylalanine (L-DOPA), a precursor of dopamine, rescues this impairment. This study establishes a simple, quick, and comparably low-cost approach that can be combined with the available genetic tools to manipulate and clarify the functions of the dopaminergic system – in *D. melanogaster* and other animals.

## Introduction

Dopamine signaling serves multiple functions, including movement initiation, sleep regulation, motivation, learning, memory extinction and forgetting (Berke, 2018; Meder et al., 2019; Oishi and Lazarus, 2017; Schultz, 2007; Yamamoto and Seto, 2014). In particular, it is crucial for conferring reinforcement signals that teach animals about the causal structure of the world (Ryvkin et al., 2018; Schultz, 2015; Waddell, 2013; Yamamoto and Vernier, 2011). This important role of dopamine is found across the animal kingdom, including the fruit fly *Drosophila melanogaster*. For this model organism, a rich genetic toolbox is available to study the functions of the dopaminergic system. Here, we suggest a complementary approach using pharmacological interventions that allow acute systemic manipulations of dopamine signaling.

Since the 1970s, both adult and larval *D. melanogaster* have been established as powerful model organisms to investigate Pavlovian conditioning, using odors as the conditioned stimulus (CS) and various types of rewarding and punishing unconditioned stimuli (US) (adults: Busto et al., 2010; McGuire et al., 2005; Perisse et al., 2013; Quinn et al., 1974; larvae: Diegelmann et al., 2013; Gerber and Stocker, 2007; Scherer et al., 2003; Thum and Gerber, 2019; Widmann et al., 2018). The genetic tools available for *D. melanogaster* allowed the genetic and neuronal mechanisms of learning and memory to be investigated, and revealed many striking similarities between the dopaminergic systems of flies and mammals, including humans (reviewed in Yamamoto and Seto, 2014). To mention but a few, flies and mammals share most genes involved in dopamine synthesis, secretion and signaling (Clark et al., 1978; Karam et al., 2019; Riemensperger et al., 2011; Yamamoto and Seto, 2014), and they also have in common the crucial role of dopaminergic neurons in reinforcement signaling (Burke et al., 2012; Liu et al., 2012; Schroll et al., 2006; Schwaerzel et al., 2003; Selcho et al., 2009; reviewed in Scaplen and Kaun, 2016). Of note is that in *D. melanogaster* different sets of dopaminergic neurons signal appetitive or aversive reinforcement, respectively, to distinct compartments of the insects’ memory center, the mushroom body, which harbors a sparse and specific representation of the olfactory environment (Diegelmann et al., 2013; Guven-Ozkan and Davis, 2014; Heisenberg, 2003; Owald and Waddell, 2015; Thum and Gerber, 2019). Due to the power, ease and elegance of the available genetic tools in *D. melanogaster*, other potentially useful techniques are often overlooked. For example, feeding or injecting drugs lacks the neuronal specificity of many transgenic tools but is a convenient way of exerting acute systemic effects. Furthermore, these approaches can be combined with genetic methods, allowing greater flexibility in manipulating the animals’ nervous system.

Interestingly, it has been demonstrated that many drugs affecting the dopamine system in mammals are also effective in flies (Nichols, 2006; Pandey and Nichols, 2011). A number of drugs that target mammalian D1 and D2 receptors have been used pharmacologically to activate and inhibit their *Drosophila* homologs in vivo (Chang et al., 2006; Srivastava et al., 2005; Yellman et al., 1997). Also, drugs that induce dopamine deficiency have been found to influence various brain functions. For example, 3-Iodo-L-tyrosine (3IY; other terms: 3-IY or 3-IT) interferes with dopamine synthesis by inhibiting the tyrosine hydroxylase enzyme (TH) that catalyzes the conversion of L-tyrosine to L-3,4-dihydroxyphenylalanine (L-DOPA), a precursor of dopamine, and it consequently reduces dopamine levels (Bainton et al., 2000; Fernandez et al., 2017; Neckameyer, 1996) **(Fig. S1).** Feeding 3IY to flies decreases activity/locomotion and increases sleep (Andretic et al., 2005; Cichewicz et al., 2017; Tomita et al., 2015; Ueno and Kume, 2014), and it alters courtship behavior (Monier et al., 2019; Neckameyer, 1998; Wicker-Thomas and Hamann, 2008). Regarding learning and memory, 3IY feeding leads to impaired visual and olfactory learning, as well as impaired long-term appetitive ethanol memory (Kaun et al., 2011; Seugnet et al., 2008; Zhang et al., 2008). Importantly, these effects of 3IY-induced dopamine deficiency can be substantially rescued by additionally feeding L-DOPA to the flies (Cichewicz et al., 2017; Monier et al., 2019; Riemensperger et al., 2011; Zhang et al., 2008).

In larvae, 3IY feeding has been used to study the developmental effects of dopamine (Neckameyer, 1996, reviewed in Verlinden, 2018) as well as the characterization of dopamine synthesis, reuptake and release (Pyakurel et al., 2018; Xiao and Venton, 2015). Furthermore, 3IY has been found to attenuate a food-odor-elicited increase in sugar feeding, an effect that was reversed by additional L-DOPA feeding (Wang et al., 2013). To our knowledge, no studies have used 3IY feeding to study learning and memory in larvae.

Here, we provide the first systematic investigation of the effects of feeding 3IY and/or L-DOPA on Pavlovian conditioning in both larval and adult *D. melanogaster*. We report detailed protocols of drug application and behavioral controls, and demonstrate the effectiveness of the approach across developmental stages (3^rd^ instar larva versus adult), valence domains (reward versus punishment), types of conditioning paradigm (absolute versus discriminatory), and types of reinforcement (natural versus optogenetically induced).

## Materials & Methods

### Animals

*Drosophila melanogaster* were raised in mass culture on standard cornmeal-molasses food and maintained at 25 °C, 60-70 % relative humidity, and a 12/12 h light/dark cycle.

For larval behavior experiments, we used 3^rd^ instar, feeding-stage wild-type Canton Special larvae of either sex, aged 4 or 5 days after egg laying, as mentioned along with the results. For adult behavior experiments, the split-GAL4 driver strain *MB320C* (detailed information can be found in the relevant database http://splitgal4.janelia.org/cgi-bin/splitgal4.cgi as well as in Aso et al., 2014), covering the PPL1-γ1pedc neurons (alternative nomenclatures: PPL1-01 and MB-MP1), was crossed to *UAS-ChR2-XXL* (Bloomington stock number: 58374, Dawydow et al., 2014) as the effector and kept in darkness throughout to avoid optogenetic activation by room light. Flies of either sex, aged 1 to 4 days after hatching, were used.

### Feeding of 3IY to larval *D. melanogaster*

A 0.5 mg/ml yeast solution was prepared from fresh baker’s yeast (common supermarket brands) diluted in tap water and stored for up to 5 days at 4 °C in a closed bottle. Samples of 2 ml yeast solution were filled into a 15 ml Falcon tube and kept for a few minutes in a warm water bath. 3-Iodo-L-tyrosine (3IY; stored at −20 °C; CAS: 70-78-0, Sigma, Steinheim, Germany) was added at a concentration of 5 mg/ml to the respective sample, if not mentioned otherwise. Notably, in contrast to earlier studies using 10 mg/ml or more (Neckameyer, 1996; Wang et al., 2013), we were not able to dissolve concentrations higher than 5 mg/ml. In some experiments, 3,4-dihydroxyphenylalanine (L-DOPA; CAS: 59-92-7, Sigma, Steinheim, Germany) was added at a concentration of 10 mg/ml, either to pure yeast solution, or to yeast solution with 5 mg/ml 3IY.

The solutions were thoroughly mixed by attaching the Falcon tubes to a shaker at high speed for approximately 60 min. Empty vials of 5 cm diameter were equipped with two layers of mesh (PET, 500 μm mesh size). Samples of the mixed yeast solution with or without additional substances were distributed onto the mesh of one vial. Larvae of the 3^rd^ instar feeding stage were collected from the fly food by adding 15 % sucrose solution (*D*-Sucrose; CAS: 57-50-1, Roth, Karlsruhe, Germany; in dH_2_O) so that the larvae floated up and could be transferred to a Petri dish filled with tap water using a tip-cut plastic pipette. After being rinsed in water, the larvae were loaded onto a filter (pluriStrainer 70 μm, pluriSelect Life Science, Leipzig, Germany) to separate them from water and small food particles, and transferred with a brush to one of the prepared vials. For yeast solutions containing different drugs and/or concentrations, different brushes were used. The larvae were left to feed on the respective yeast solution for 24 or 4 hours at 25 °C and 60-70 % relative humidity. The desired number of larvae were collected with a brush, briefly rinsed in water, and afterwards used in the respective experiment.

### Larval behavior

#### Odor-fructose associative learning

Experiments for appetitive odor-fructose associative memory (Saumweber et al., 2011; Scherer et al., 2003) were performed using a one-odor, single-training-trial protocol described in Weiglein et al. (2019). For example, two custom-made Teflon containers of 5 mm diameter were filled with 10 μl of odor substance (*n*-amylacetate, AM; CAS: 628-63-7, Merck, Darmstadt, Germany; diluted 1:20 in paraffin oil; CAS: 8042-47-5, AppliChem, Darmstadt, Germany) and closed with lids perforated with 5-10 holes, each of approximately 0.5 mm diameter. These odor containers were located on opposite sides of a Petri dish (9 cm inner diameter; Nr. 82.1472 Sarstedt, Nümbrecht, Germany) filled with 1 % agarose solution (electrophoresis grade; CAS: 9012-36-6, Roth, Karlsruhe, Germany) and additionally containing fructose (FRU; 2 M; purity 99 %; CAS: 57-48-7 Roth, Karlsruhe, Germany) as a taste reward (+). Cohorts of approximately 30 larvae were placed at the center of the Petri dish and allowed to move about the Petri dish for 2.5 min. Subsequently, they were transferred with a brush to a fresh Petri dish that was filled with plain, tasteless agarose and equipped with two empty Teflon containers (EM). For each cohort trained in such a paired way (paired training; AM+/EM), a second cohort of larvae received the odor unpaired from the fructose reward (unpaired training; EM+/AM). In half of the cases the order of sequence was reversed (EM/AM+, AM/EM+, respectively).

After one training trial, the larvae were transferred to a fresh, tasteless test Petri dish with AM on one side and an EM container on the opposite side. The larvae were left to distribute for 3 min and then counted to evaluate their preference for AM. The number of larvae (#) on the AM side, on the EM side, and in a 10-mm wide middle zone was counted. Larvae crawling up the sidewalls of the Petri dish were counted for the respective side, whereas larvae on the lid were excluded from the analysis. A preference index (AM PREF) was calculated:

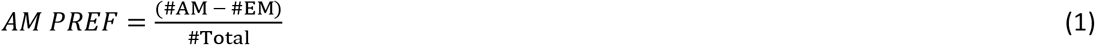

AM PREF values range from +1 to −1, with positive values indicating odor preference and negative values indicating avoidance of AM.

From the AM PREF scores after paired and unpaired training, a performance index (PI) was calculated as follows:

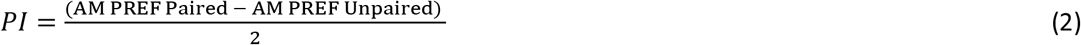

Performance indices range from +1 to −1. Positive PIs indicate appetitive associative memory; negative values indicate aversive associative memory.

#### Innate odor preference tests

Cohorts of approx. 20–30 experimentally naïve larvae were collected, briefly washed in tap water, and placed onto a Petri dish with an AM container on one side and an empty container on the other side. After 3 minutes, the odor preference was determined as detailed in equation (1).

#### Innate fructose preference tests

Split Petri dishes were prepared freshly approx. 4 hours before the experiment, following the procedures described in König et al. (2014) such that one half of the Petri dish (9 cm diameter) was filled with agarose with 2 M fructose (FRU), and the other half with plain agarose. Approx. 20–30 larvae were collected, rinsed in tap water, and placed onto the center of a split Petri dish. After 3 min, the number of larvae (#) on the fructose side, on the pure agarose side, and in a 10-mm wide middle zone was counted. Fructose preference was calculated as follows:

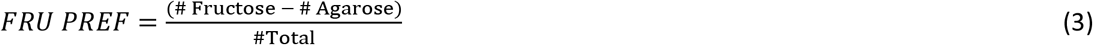

FRU PREF scores range from +1 to −1, with positive values indicating fructose preference and negative values indicating avoidance.

#### Analyses of locomotion

Cohorts of approximately 20 larvae were placed on an empty, plain-agarose-filled Petri dish without odor or reward. For three minutes, they were video-recorded while they freely moved in the dish. The videos were analyzed offline using custom-made tracking software described in Paisios et al. (2017). In brief, larvae alternately perform relatively straight forward locomotion, called runs, and lateral head movements, called head casts (HCs) that are often followed by changes in direction. This leads to a typical zig-zagging pattern of locomotion (Gershow et al., 2012; Gomez-Marin and Louis, 2014; Gomez-Marin et al., 2011). As described in detail by Paisios et al. (2017), an HC was detected whenever the angular velocity of a vector through the animal’s head exceeded a threshold of 35 °/s and ended as soon as that angular velocity dropped below the threshold again. The time during which an animal was not head-casting was regarded as a run, deducting 1.5 seconds before and after an HC to exclude the decelerating and accelerating phases that usually happen before and after an HC, respectively. Three aspects of behavior were analyzed:

- the run speed was determined as the average speed (mm/s) of the larval midpoint during runs;
- the rate of HCs was determined as the number of HCs per second (HC/s);
- the size of HCs was determined by the HC angle. Accordingly, the animal’s bending angle as the angle between vectors through the head and tail was determined before and after an HC. Then, the HC angle was calculated as the difference between the animal’s bending angle after an HC and the bending angle before an HC. For a detailed description, see Paisios et al. (2017).

To analyze the HC behavior in more detail, we determined the HC rate and HC angle separately for small and large HCs. The discriminatory threshold for large HCs of an HC angle > 20° was based on previous studies (Paisios et al., 2017; Schleyer et al., 2015; Thane et al., 2019).

### Feeding of 3IY to adult *D. melanogaster*

For 3IY feeding in adult flies, a 5 % sucrose solution (CAS: 57-50-1, Hartenstein, Würzburg, Germany) was prepared. This solution was either used pure, or mixed with 5 mg/ml 3IY, or with 10 mg/ml L-DOPA, or with both, in an analogous manner to that described above for the larval case. Hatched adults of the genotype *MB320C;ChR2-XXL* were collected in fresh food vials and kept under the normal culture conditions mentioned above, at least overnight and at most until 4 days after hatching. Flies were transferred to new vials containing a tissue (Fripa, Düren, Germany) soaked with 1.8 ml of sucrose solution that either did or did not contain 3IY and/or L-DOPA, as mentioned in the results section. After 40-48 h under otherwise normal culture conditions, the flies were trained and / or tested *en masse*, in cohorts of ∼100 in the respective paradigm.

### Adult behavior

#### Odor-PPL1-γ1pedc associative learning

For the memory assays, we followed the procedures described in König et al. (2018), unless mentioned otherwise. Flies were loaded into a small transparent tube in a custom-made set-up (CON-ELEKTRONIK, Greussenheim, Germany) and were trained and tested at 23–25°C and 60–80% relative humidity. Training was performed in dimmed red light, which is largely invisible to flies and does not stimulate the ChR2-XXl effector; testing was performed in darkness. For the application of blue light, a 2.5 cm-diameter and 4.5 cm-length hollow tube with 24 LEDs mounted on the inner surface was placed around the transparent training tubes harboring the flies. As odorants, 50 μl benzaldehyde (BA) and 250 μl 3-octanol (OCT) (CAS 100-52-7, 589-98-0; both from Fluka, Steinheim, Germany) were applied to 1 cm-deep Teflon containers of 5 and 14 mm diameter, respectively. From these, odor-loaded air was shunted into the permanent air stream flowing through the apparatus. During training, the flies were presented with both odors for 1 min with a 3 min resting interval in between, but only one of the odors was paired with 1 min of blue light (465 nm) for optogenetic activation of PPL1-γ1pedc, whereas the other odor was presented alone (either BA-paired or OCT-paired training, respectively). In half of the cases training started with the odor paired with light (CS+); in the other half training started with the odor without light activation (CS-; for details see electronic supplement figure S1B of König et al., 2019). For the subsequent test, the flies were given a 3 min accommodation period, after which they were transferred to the T-maze-like choice point. The test configuration between the two odors used during training was prepared and balanced so that either BA or OCT were present at front vs rear position over the course of all experiments. After 2 min testing time, the arms of the maze were closed and the flies on each side were counted to calculate a benzaldehyde preference index (BA PREF):

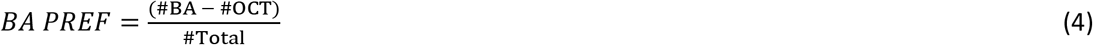

Thus, positive scores indicate preference for BA and negative scores preference for OCT. From the BA PREF scores of two independently trained fly groups after BA-paired and OCT-paired training, a performance index (PI) was calculated as follows:

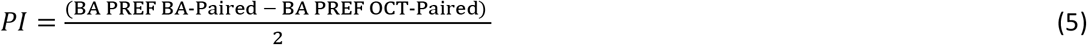

Positive performance indices thus reflect appetitive associative memory, negative values aversive associative memory.

#### Innate odor preference tests

Cohorts of ∼ 50 flies were loaded into the setup. After a 5 min resting interval, they were transferred to the choice point of a T-maze between an arm equipped with either BA or OCT, and an arm with an empty Teflon container, and allowed to distribute for 2 min. A Preference was calculated as:

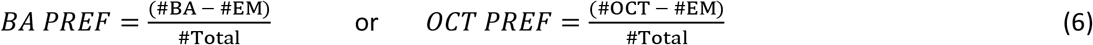

Each data point in Fig. 4C and D represents the mean value of 2 runs tested with the odor in the front or rear T-Maze position.

### Statistics

Two-tailed, non-parametric statistics were used throughout to analyze the behavioral data. For comparisons of a group’s scores with chance levels (zero), one-sample sign tests (OSS) were applied. To compare across multiple independent groups, Kruskal-Wallis tests (KW) with subsequent pair-wise Mann-Whitney U-tests (MWU) were used (Statistica 13, StatSoft Inc, Tulsa, USA). To ensure a within-experiment error rate below 5%, a Bonferroni-Holm (BH) correction for multiple comparisons was employed (Holm, 1979). Sample sizes (biological replications) were estimated based on previous studies with small to medium effect sizes (König et al., 2018; Weiglein et al., 2019). None of the specific experiments reported here had previously been performed in our laboratory, although the basic behavioral paradigms are regularly used. Experimenters were blind to treatment condition during the experiments with larvae, and during the fly counting for the experiments with adults. Data are presented as box plots showing the median as the middle line, the 25 and 75 % quantiles as box boundaries, and the 10 and 90 % quantiles as whiskers. All data from behavioral experiments are documented in the Supplemental Data file ‘Table S1’.

## Results

### Feeding 3IY for 24 hours induces broad behavioral impairments in larvae

We first investigated the effects of 3IY feeding on *D. melanogaster* larvae. In an approach modified from Neckameyer (1996), cohorts of 4-day-old larvae were placed on a PET mesh soaked with a yeast solution mixed with 3IY at the indicated concentrations, or without 3IY. After 24 hours, the larvae underwent a single-trial Pavlovian training with odor and sugar reward, following established protocols (Michels et al., 2017; Saumweber et al., 2011; Scherer et al., 2003; Weiglein et al., 2019): one cohort of larvae was trained by a paired presentation of odor and reward, whereas a second cohort was trained reciprocally, i.e. with separated, unpaired presentations of odor and reward. In control larvae that were kept on a yeast solution without 3IY, an appetitive associative memory was revealed by higher odor preferences after paired than after unpaired training in a subsequent test (**Fig. S2A**), indicated by positive performance index (PI) scores (**Fig. 1A**, left-most box plot). When we performed the same learning experiment with larvae fed with various concentrations of 3IY, we observed decreased memory scores with increased 3IY concentrations. Significantly reduced scores were found for a concentration of 5 mg/ml (**Fig. 1A** and **S2A**), a result we replicated in an independent experiment (**Fig. 1B** and **S2B**). However, we noticed that many larvae had died due to the treatment, and the cuticle of many of the surviving animals was darkened (not shown). We therefore wondered whether the treatment may generally impair behavioral faculties. Indeed, innate odor preference was found to be impaired in 3IY-fed larvae (**Fig. 1C**). This prompted us to test their basic locomotion on an empty, tasteless Petri dish without odor or sugar, and to analyze their behavior using custom-made analysis software (Paisios et al., 2017). Typically, larvae move by relatively straight runs, interrupted by turning maneuvers indicated by lateral head movements called head casts (HC) (**Fig. S2C**) (Gershow et al., 2012; Gomez-Marin and Louis, 2014; Gomez-Marin et al., 2011; Paisios et al., 2017; Thane et al., 2019). Analysis of these parameters of locomotion revealed that the animals’ run speed was unchanged by 3IY feeding (**Fig. 1D**). However, the larvae fed with 3IY systematically performed fewer and larger HCs than control animals (**Fig. 1E-F, Fig. S2D-G**).

**Figure 1:**
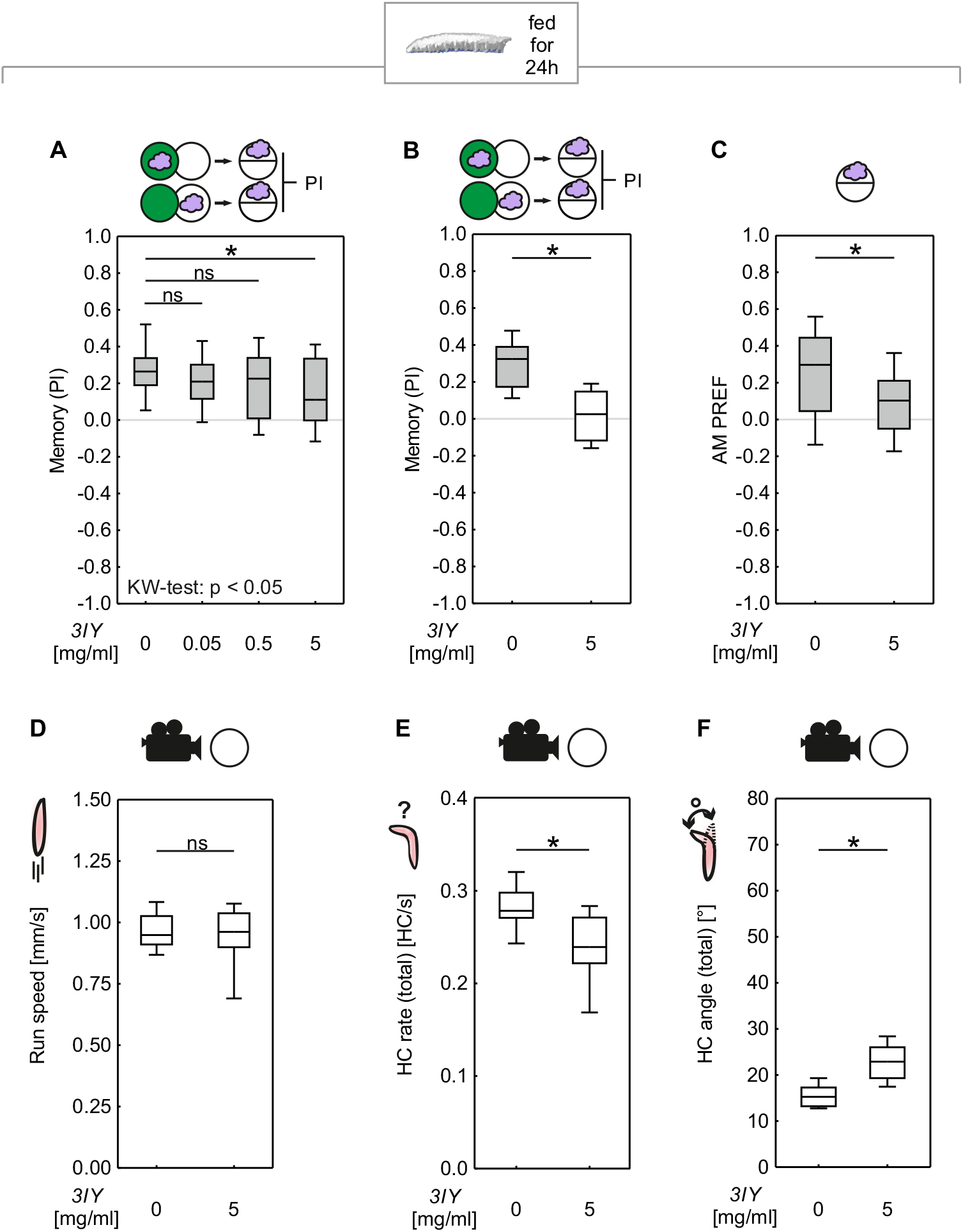
Feeding 3IY to *D. melanogaster* larvae for 24h broadly impairs behavior. **(A)** Cohorts of larvae were trained by either paired or unpaired presentations of an odor (purple cloud) and sugar (green circle), and subsequently tested for odor preference. Feeding different concentrations of 3IY for 24h led to memory impairment (KW: *H* = 8.44, d.f. = 3, *P =* 0.378; OSS from left to right: *P* < 0.0001; *P* < 0.0001; *P =* 0.0019; *P =* 0.0039; N = 36 each), with a significant reduction compared to the control only in the group with the highest tested concentration of 5mg/ml 3IY (MWU: *U* = 405.00, *P =*0.0063). All other tested concentrations did not affect memory scores compared to the control group (MWU: 0 vs. 0.05 mg/ml 3IY: *U* = 533.50, *P =* 0.1992; 0 vs. 0.5 mg/ml 3IY: *U* = 518.00, *P =* 0.1447). **(B)** As seen in (A), larvae fed with 5mg/ml 3IY showed impaired memory (MWU: *U* = 19.00, *P =* 0.0024; OSSs from left to right: *P =* 0.0005; *P =* 0.7744; N = 12 each) in an independent repetition. **(C)** An innate preference test revealed lower preference for the tested odor in the group fed with 5 mg/ml 3IY compared to the control group (MWU: *U* = 852.00, *P =* 0.0061; OSSs from left to right: *P* < 0.0001; *P =* 0.0066; N = 50 each). **(D)** Offline analysis of larval behavior revealed no difference in run speed between control larvae and larvae fed with 5 mg/ml 3IY (MWU: *U* = 192.00, *P =* 0.8392, N = 20 each). Regarding head casts, larvae fed with 5 mg/ml 3IY compared to control larvae showed **(E)** fewer head casts (MWU: *U* = 76.00, *P =* 0.0008, N = 20 each) but **(F)** made larger head casts (MWU: *U* = 28.00, *P* < 0.0001, N = 20 each). Grey boxes reflect memories relative to chance levels (PI = 0) significant at *P* < 0.05 in OSS-tests with Bonferroni-Holm correction. KW-tests are indicated within the figure. Asterisks and ns above horizontal lines reflect significance or lack thereof in MWU-tests. Box plots represent the median as the midline, 25 and 75% as the box boundaries, and 10 and 90% as the whiskers. See Figure S2 for preference scores underlying the PIs and detailed head cast analysis.

Thus, feeding the larvae with 5 mg/ml 3IY for 24 hours seemed to impair their basic behavioral faculties, suggesting that the reduced memory scores that we observed after the treatment might be secondary to such general impairment. Therefore, we next sought to reduce the ‘side effects’ of 3IY feeding.

### Feeding 3IY for 4 hours specifically impairs associative learning in larvae

Given the reported role of dopamine and the TH enzyme in development and cuticle formation (Friggi-Grelin et al., 2003; Hsouna et al., 2007; Neckameyer, 1996; Neckameyer and White, 1993; reviewed in Verlinden, 2018), the timing of 3IY feeding is likely to have an impact. In order to minimize developmental effects, it seems desirable to apply 3IY as late as possible in the larval life cycle (and yet early enough to be able to finish the experiment before the larvae start to pupate). We therefore reduced the duration of 3IY feeding to four hours, which allowed for the feeding of 3IY to 5-day-old animals. After this shortened feeding protocol too, memory scores were reduced compared to controls (**Fig. 2A** and **S3A**). Critically, the animals’ basic behavioral faculties turned out to be intact: no impairment in innate odor preference (**Fig. 2B**) or sugar preference (**Fig. 2C**) was detectable. Thus, the shortened feeding of 3IY specifically impaired associative memory without impairing task-relevant behavioral faculties (nor did we observe any dead or darkened larvae; not shown). A more detailed analysis of locomotion revealed only a very mild increase in the HC rate but otherwise no differences with respect to controls (**Fig. 2D-F**, for a more detailed analysis, see **Fig. S3B-E**).

**Figure 2:**
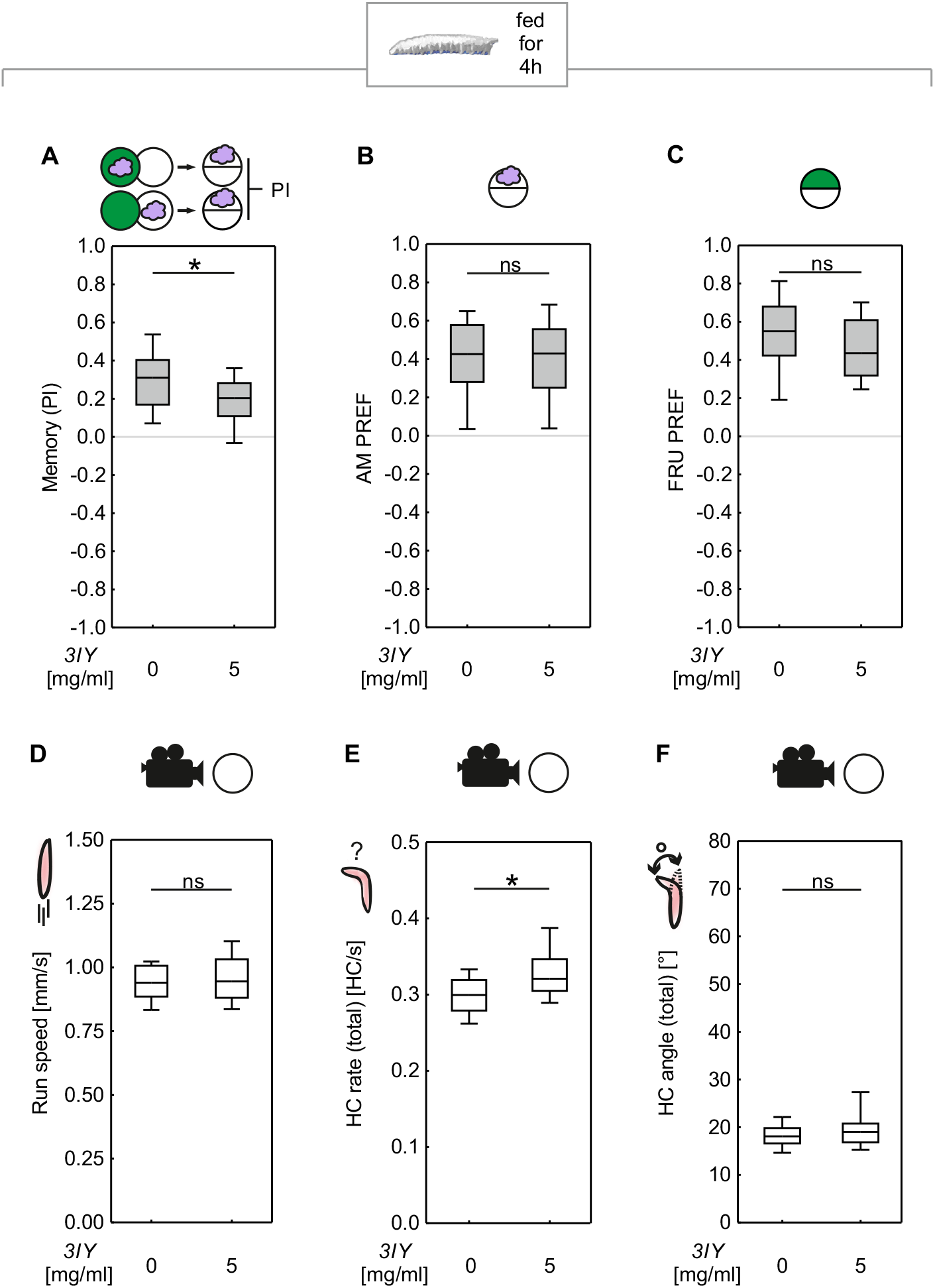
Feeding 3IY to *D. melanogaster* larvae for 4h impairs memory but leaves innate behavior intact. **(A)** Larvae fed with 5 mg/ml 3IY for 4h showed impaired memory compared to the control group (MWU: *U* = 125.00, *P =* 0.0439; OSS from left to right: *P* < 0.0001; *P =* 0.0004; N = 20 each). Innate preference for the **(B)** odor (MWU: *U* = 230.50, *P =* 0.7963; OSS: *P* < 0.0001 each; N = 22 each) or **(C)** FRU (MWU: *U* = 344.00, *P =* 0.1188; OSS: *P* < 0.000 each; N = 30 each) was not affected. Video-tracking of the larvae revealed **(D)** no difference in run speed (MWU: *U* = 254.00, *P =* 0.4897, N = 24 each), **(E)** a slight increase in HC rate for larvae fed with 5mg/ml 3IY (MWU: *U* = 174.00, *P =* 0.0193, N = 24 each), and **(F)** no difference in HC angles (MWU: *U* =247.00, *P =* 0.4037; N = 24 each). See Figure S3 for preference scores underlying the PIs and detailed analysis of head casts. For further details, see Figure 1.

We next tried to rescue the effect of 3IY on the TH enzyme by additionally feeding the animals with L-DOPA (**Fig. S1**). To this end, we fed animals either with plain yeast solution (control), or 5 mg/ml 3IY, or with both 5 mg/ml 3IY and 10 mg/ml L-DOPA. The memory scores were impaired in larvae fed with 3IY alone (**Figure 3A**), replicating the results from Figure 2A. These reduced memory scores were restored to control levels by additionally feeding L-DOPA to the larvae (**Fig. 3A** and **S4A**). Innate odor and sugar preferences were not affected by either 3IY or combined 3IY and L-DOPA feeding, confirming that both effects were specific for associative learning (**Fig. 3B-C**). Importantly, while a repetition of the experiment from Figure 3A replicated the finding that L-DOPA feeding can restore memory scores upon 3IY treatment, we also showed that the feeding of L-DOPA alone did not increase memory scores (**Fig. 3D** and **S4B**).

**Figure 3:**
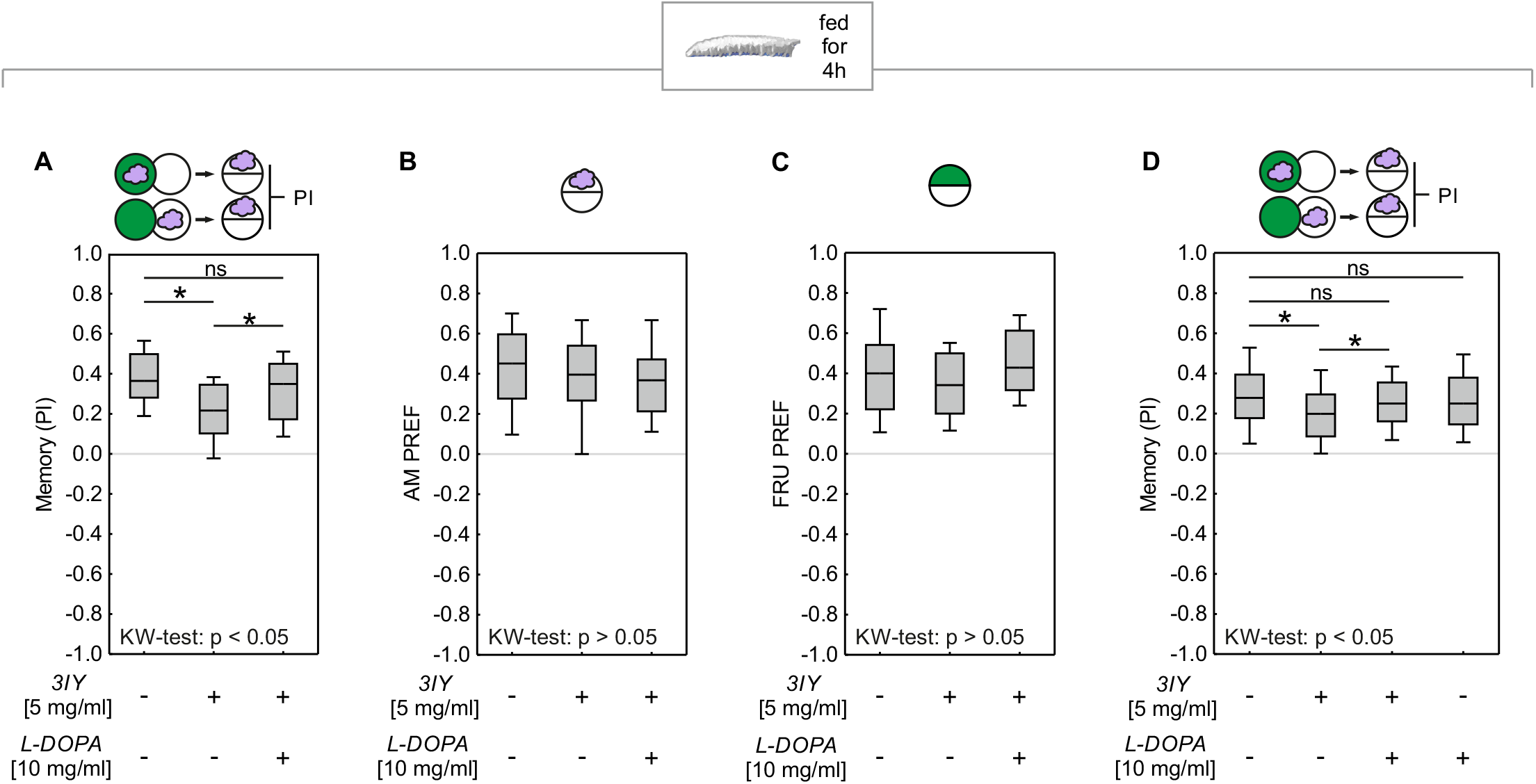
Memory impairment in *Drosophila* larvae due to 3IY can be rescued by additionally feeding L-DOPA. **(A)** Feeding L-DOPA in addition to 3IY rescued the memory impairment (KW: *H* = 10.69, d.f. = 2, *P =* 0.0048; OSS from left to right: *P* < 0.0001; *P =* 0.0005; *P* < 0.0001; N = 26 each). Feeding 5 mg/ml 3IY alone for 4h impaired memory (MWU: *U* = 165.50, *P =* 0.0016) whereas additionally feeding 10 mg/ml L-DOPA rescued memory impairment (MWU: *U* = 215.00, *P =* 0.0250) and led to memory scores comparable to the control group (MWU: *U* = 286.00, *P =* 0.3459). Feeding either drug did not affect innate approach to **(B)** odor (KW: *H* = 2.02, d.f. = 2, *P =* 0.3650; OSS from left to right: *P* < 0.0001; *P* < 0.0001; *P* < 0.0001; N = 28 each) or **(C)** FRU (KW: *H* = 2.42, d.f. = 2, *P =* 0.2977; OSSs from left to right: *P* < 0.0001; *P* < 0.0001; *P* < 0.0001; 24 each). **(D)** As shown in (A), feeding 3IY impaired memory scores and this impairment was rescued by additional L-DOPA feeding (KW: *H* = 14.06, d.f. = 3, *P =* 0.0028; OSS: *P* < 0.0001 each; N = 96 each; MWU: no drug vs. 3IY alone: *U* = 3262.50, *P =* 0.0005; no drug vs. 3IY + L-DOPA: *U* = 4084.50, *P =* 0.1743; 3IY alone vs. 3IY + L-DOPA: *U* = 3673.00, *P =* 0.0152). Feeding L-DOPA alone had no effect on memory scores (MWU: no drug vs. L-DOPA alone: *U* = 4246.50, *P =* 0.3484). See Figure S4 for preference scores underlying the PIs. For further details, see Figure 1.

### Feeding of 3IY specifically impairs associative learning in adults

After demonstrating the effectiveness of 3IY feeding on associative learning about natural sugar rewards in larvae, we sought to study how broadly applicable the 3IY approach might be. Therefore, we applied it to a very different learning paradigm, by using adult flies instead of larvae; a two-odor discrimination paradigm instead of a one-odor, non-discriminatory paradigm; and an optogenetic punishment instead of a natural taste reward (**Fig. 4**). Specifically, we expressed the blue-light-gated cation channel channelrhodopsin-2-XXL as the optogenetic effector (*ChR2-XXL*; Dawydow et al., 2014) in a single dopaminergic neuron per brain hemisphere, called PPL1-γ1pedc (alternative nomenclatures: PPL1-01 and MB-MP1), as covered by the Split-GAL4 driver strain *MB320C* (Aso et al., 2014). This neuron, when optogenetically activated, carries an internal punishment signal sufficient to establish an aversive associative memory when paired with an odor (Aso and Rubin, 2016; Hige et al., 2015; König et al., 2018) (**Fig. 4A, left-most box plot**). Upon feeding 3IY for 48 hours before training, memory scores were decreased, an effect that was restored by L-DOPA feeding (**Fig. 4A and S5A**). The effect of 3IY in reducing memory scores increased with increasing 3IY concentrations (**Fig. 4B** and **S5B**). 3IY feeding left innate odor preference to either odor unaffected (**Fig. 4C-D**) and feeding L-DOPA alone did not increase memory scores (**Fig. 4E and S5C**). Thus, feeding 3IY specifically impaired associative learning in adult flies, but kept their task-relevant behavioral capacities intact.

**Figure 4:**
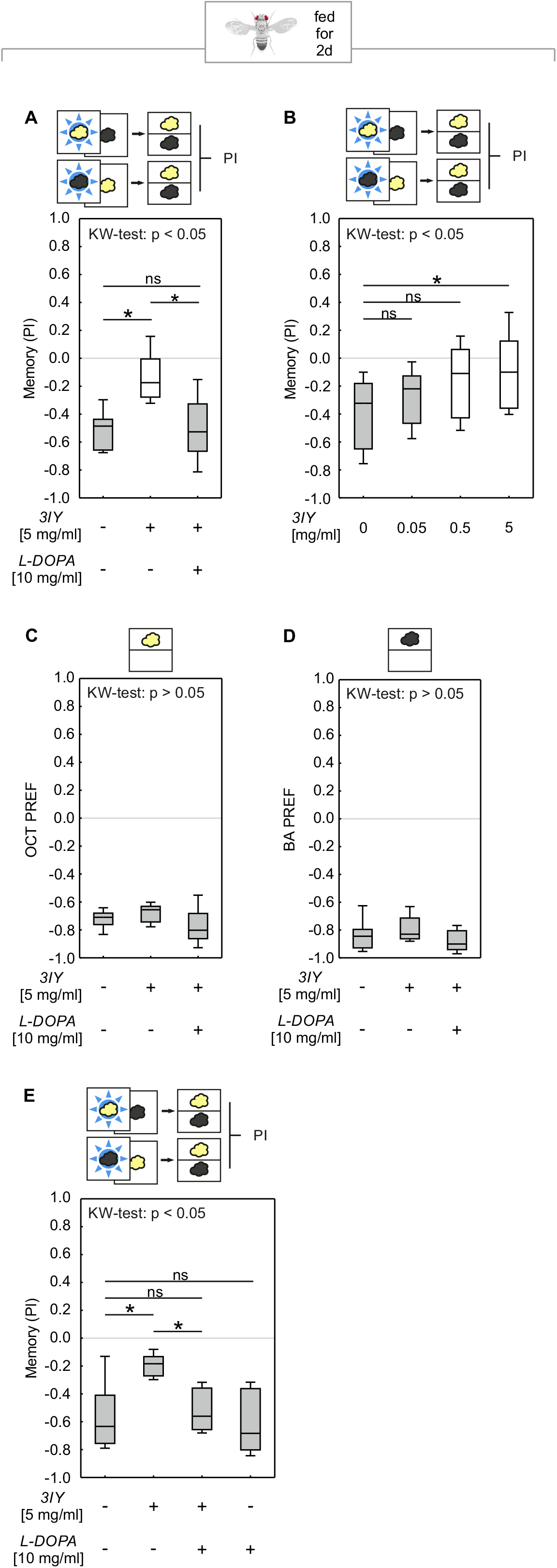
Feeding 3IY to adult *D. melanogaster* impairs optogenetically induced memory but leaves innate behavior intact. **(A)** Cohorts of flies were trained by pairing one of two odors (yellow / black cloud) with optogenetic activation of PPL1-γ1pedc (blue star), and subsequently tested for their choice between the two odors. 3IY feeding led to an impaired performance index compared to the control group (MWU: *U* = 2.00, *P =* 0.0019). Additional L-DOPA feeding (KW: *H* = 11.89, d.f. = 2, *P =* 0.0026; OSS from left to right: *P =* 0.0078; *P =* 0.2891; *P =* 0.0078; N = 8 each) rescued this impairment of memory scores (MWU: 3IY alone vs 3IY+L-DOPA: *U* = 6.00, *P =* 0.0074) to the control level (MWU: no drug vs 3IY+L-DOPA: *U* = 29.00, *P =* 0.7929). **(B)** 3IY concentrations significantly influenced PI values (KW: *H* = 11.08, d.f. = 3, *P =* 0.0113; OSS from left to right: *P =* 0.0005; *P =* 0.0018; *P =* 0.2101; *P =* 1.0; N = 16, 14, 16, 15). The highest concentration of 5 mg/ml 3IY significantly reduced memory compared to the control group (MWU: *U* = 47.00, *P =* 0.0042). All other tested concentrations of 3IY had no significant effect with the given sample sizes (MWU: 0 vs. 0.05 mg/ml 3IY: U = 83.00, *P =* 0.2361; 0 vs. 0.5 mg/ml 3IY: U = 72.00, *P =* 0.0365). **(C-D)** Innate odor avoidance of **(C)** OCT and **(D)** BA was not affected by 3IY and / or L-DOPA feeding (KW: OCT: *H* = 4.42, d.f. = 2, *P =* 0.1097; OSS from left to right: *P* < 0.0001; *P* < 0.0001; *P* < 0.0001; N = 12 each) (KW: BA: *H* = 2.71, d.f. = 2, *P =* 0.2575; OSS from left to right: *P* < 0.0001; *P* < 0.0001; *P* < 0.0001; N = 12 each) **(E)** In a repetition of the experiment shown in (A), feeding L-DOPA in addition to 3IY rescued the 3IY-induced memory impairment (KW: *H* = 14.68, d.f. = 3, *P =* 0.0021; OSS: *P* < 0.0001 each; N = 13, 12, 11, 12; MWU: no drug vs. 3IY alone: *U* = 29.00, *P =* 0.0083; no drug vs. 3IY + L-DOPA: *U* = 59.00, *P =* 0.4869; 3IY alone vs. 3IY + L-DOPA: *U* = 12.00, *P* = 0.0009). Importantly, L-DOPA alone had no effect on the memory scores (MWU: no drug vs. L-DOPA alone: *U* = 68.00, *P =* 0.6053). See Figure S5 for preference scores underlying the PIs. For further details, see Figure 1.

## Discussion

The present study demonstrates that both in larval and adult *D. melanogaster*, and in two very different kinds of task, feeding 3IY can specifically impair associative learning while innate behavior remains intact. In either case, the observed memory impairment was rescued by feeding L-DOPA, suggesting that the 3IY-impairment was indeed caused by an inhibition of the tyrosine hydroxylase enzyme that catalyzes the synthesis of L-DOPA. These results confirm previous studies that had used genetic tools to demonstrate the critical role of dopamine for associative learning in *D. melanogaster* (Himmelreich et al., 2017; Kim et al., 2007; Riemensperger et al., 2011; Rohwedder et al., 2016; Selcho et al., 2009).

This study provides an approach to manipulating the dopaminergic system of *D. melanogaster* independently of genetic tools. It is easy to apply, quick, and comparably cheap. It does not require generating new fly strains, but can be easily combined with the use of already available genetic tools that would manipulate the dopaminergic system in other ways, or that target different types of processes (**Fig. 4**). Also, the effects of the drugs can be titrated relatively conveniently by adjusting the concentration and the duration of feeding. This makes it possible to find a trade-off between maximizing the intended effect on learning and memory and minimizing developmental side effects. Furthermore, drugs such as 3IY, with comparable effects in different organisms, allow for elegant translational research across different species. Finally, it may actually be an advantage that systemic pharmacological approaches affect the target process in the complete organism at once. For example, using the genetic driver strain TH-Gal4, which was widely believed to cover all dopaminergic neurons, Schwaerzel et al. (2003) suggested that dopaminergic neurons were responsible only for punishment, but not reward signaling (see also Schroll et al., 2006, regarding larvae). This needed to be reconsidered when it turned out that the driver strain in question missed a crucial cluster of dopaminergic neurons that do indeed signal reward (Burke et al., 2012; Liu et al., 2012; larvae: Rohwedder et al., 2016). A systemic pharmacological approach could have made this discovery possible many years earlier.

Taken together, the pharmacological approach introduced here adds to the toolbox for *Drosophila* neuroscience and will hopefully help us to understand the functions of the dopaminergic system and the brain in general – in *D. melanogaster* and other animals.

## Acknowledgements

Discussions with B. Gerber and P. Stevenson, as well as technical assistance by A. Ciuraszkiewicz-Wojciech, B. Kracht, T. Niewalda and M. Thane, are gratefully acknowledged. We thank R.D.V. Glasgow (Zaragoza, Spain) for language editing.

## Author contributions

CK, JT, MS, AW, NT and NM conceived and planned the study; CK, JT, AW, NT, NM and FA performed the experiments; JT, CK and MS analyzed the data; MS, JT, CK and AW wrote the manuscript with input of all authors.

---

**Figure S1:**
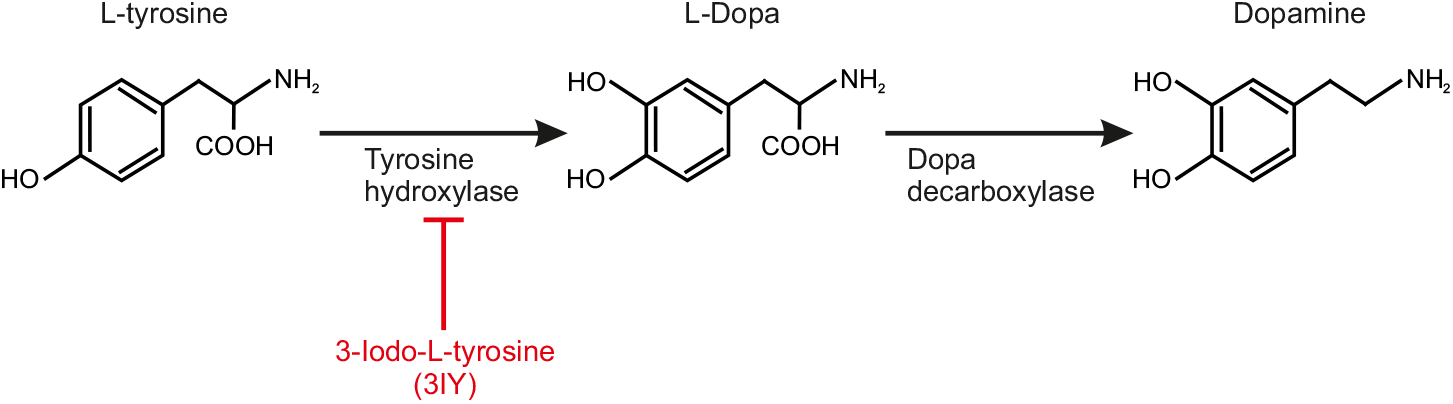
Dopamine synthesis. Dopamine is synthesized via two enzymatic steps. In the first step the amino acid L-tyrosine is converted into L-3,4-dihydroxyphenylalanine (L-DOPA) via tyrosine hydroxylase (TH). In the second step, L-DOPA is converted to dopamine via dopa decarboxylase (DDC). In red, the inhibition of the TH enzyme by 3-Iodo-L-tyrosine (3IY) is shown.

**Figure S2:**
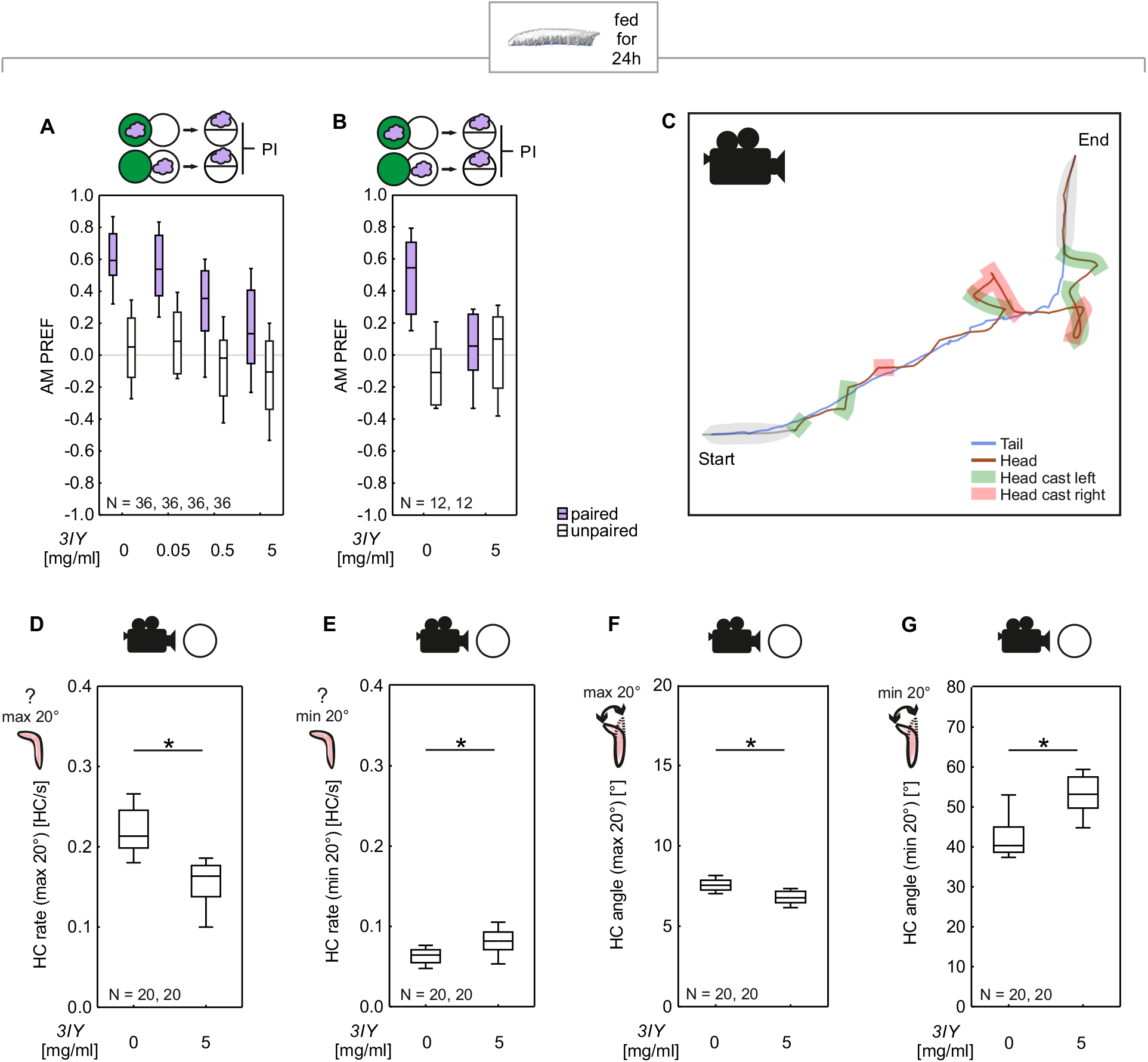
Preference and tracking data underlying the results shown in Figure 1. **(A, B)** Preference data refer to the Performance Indices shown in Figure 1A and B, respectively. Purple boxes represent odor preference after paired training (AM+), white boxes after unpaired training (EM+). **(C)** Example of a track from the video-recording of a single larva showing relatively straight runs interrupted by lateral head movements (head cast, HC). **(D, E)** HC rate for small and large HCs, respectively, classified by a HC angle smaller or greater than 20°. This classification as well as the calculation of the HC angle is based on Paisios et al. (2017). The HC rate was decreased for small HCs **(D)** (MWU: U = 29.00, *P* < 0.0001, N = 20 each) and increased for large HCs **(E)** (MWU: U = 85.00, *P =* 0.0019, N = 20 each) for larvae fed with 3IY for 24 h. **(F, G)** HC angles classified by small and large HCs. The average HC angle was decreased for small HCs **(F)** (MWU: U = 47.00, *P* < 0.0001, N = 20 each) and increased for large HCs **(G)** (MWU: U = 46.00, *P* < 0.0001, N = 20 each) for larvae fed with 3IY for 24h. Asterisks above horizontal lines reflect significance in MWU-tests. Box plots represent the median as the midline, 25 and 75% as the box boundaries, and 10 and 90% as the whiskers. Sample sizes are indicated within the figure.

**Figure S3:**
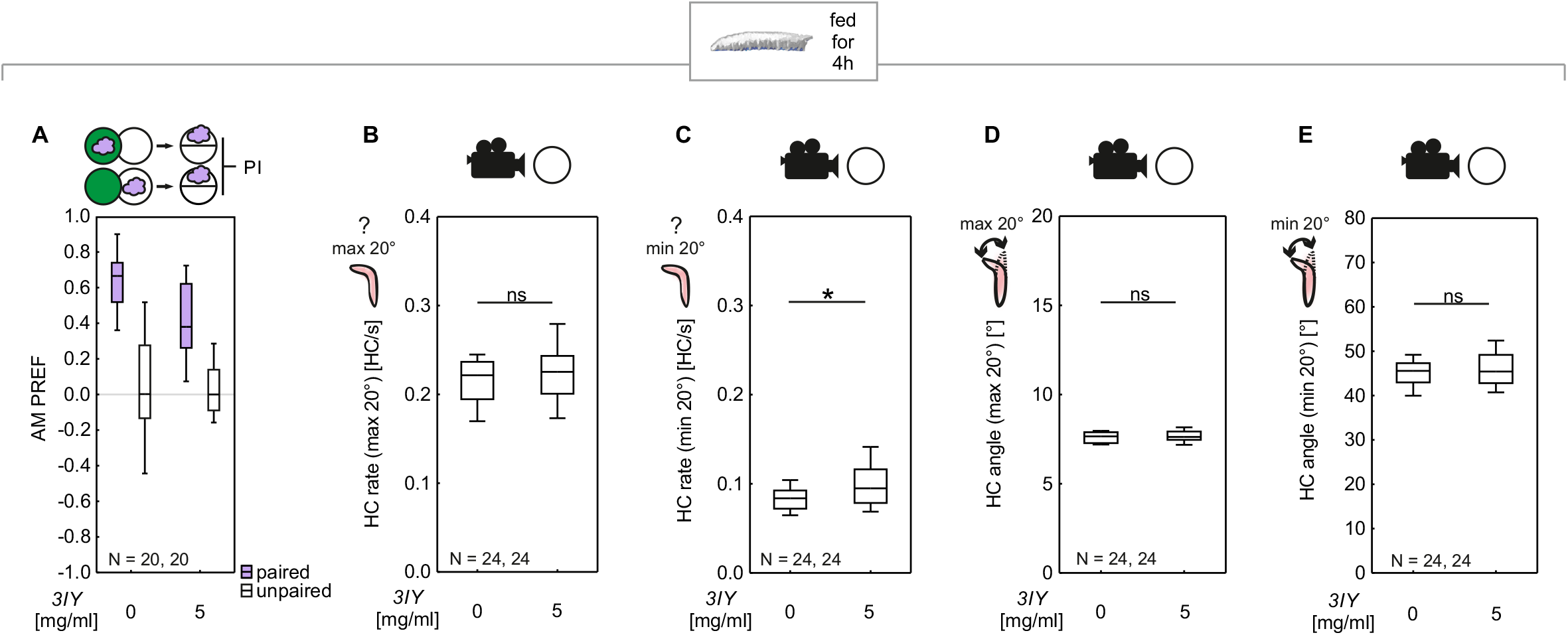
Preference and tracking data underlying the results shown in Figure 2. **(A)** Preference data refer to the Performance Indices shown in Figure 2A. Purple boxes represent odor preference after paired training (AM+), white boxes after unpaired training (EM+). **(B, C)** HC rate classified by small and large HCs. The HC rate for small HCs was unaffected by 3IY feeding **(B)** (MWU: U = 231.00, *P =* 0.2440, N = 24 each), but slightly increased for large HCs **(C)** (MWU: U = 186.00, *P =* 0.0364, N = 24 each). Comparing these results with those after 24 h feeding (Fig. S2D-E), it seems that both 4 h and 24 h feeding of 3IY slightly increased the rate of large HCs, but only 24 h feeding decreased the rate of small HCs in addition. This resulted in a decrease in total HC rate after 24 h feeding on 3IY as seen in Fig. 1E, and in an increase in total HC rate after 4 h feeding on 3IY as shown in Fig. 2E. **(D, E)** HC angles classified by small and large HCs. No significant effect of 3IY feeding was observed (MWU: HC angle max 20°: U = 256.00, *P =* 0.5160, N = 24 each; HC angle min 20°: U = 268.00, *P =* 0.6876, N = 24 each). Asterisks and ns above horizontal lines reflect significance or lack thereof in MWU-tests. Box plots represent the median as the midline, 25 and 75% as the box boundaries, and 10 and 90% as the whiskers. Sample sizes are indicated within the figure.

**Figure S4:**
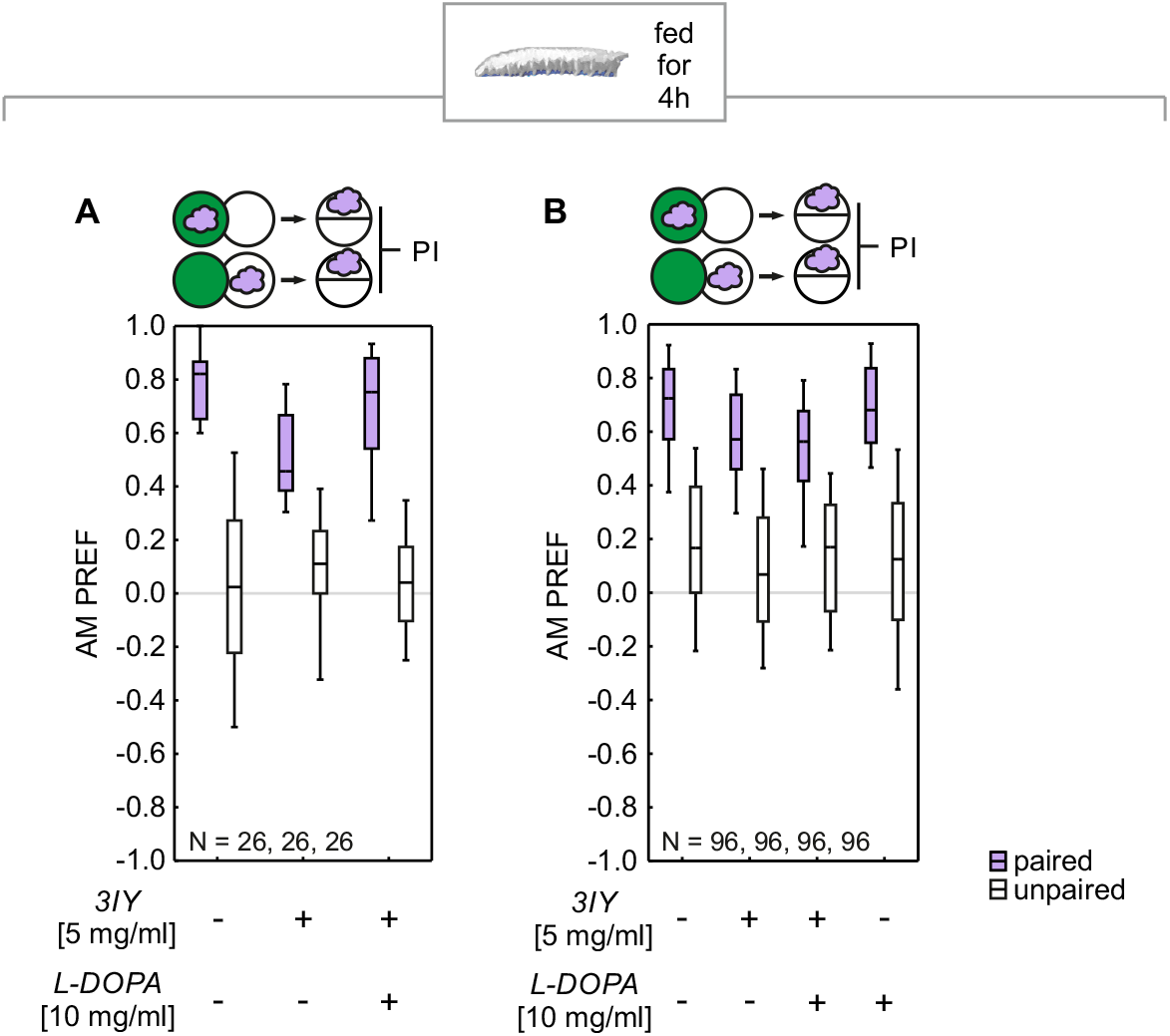
Preferences underlying the Performance Indices in Figure 3. **(A, B)** Preference data refer to the Performance Indices shown in Figure 3A and D, respectively. Purple boxes represent odor preference after paired training (AM+), white boxes after unpaired training (EM+). Box plots represent the median as the midline, 25 and 75% as the box boundaries, and 10 and 90% as the whiskers. Sample sizes are indicated within the figure.

**Figure S5:**
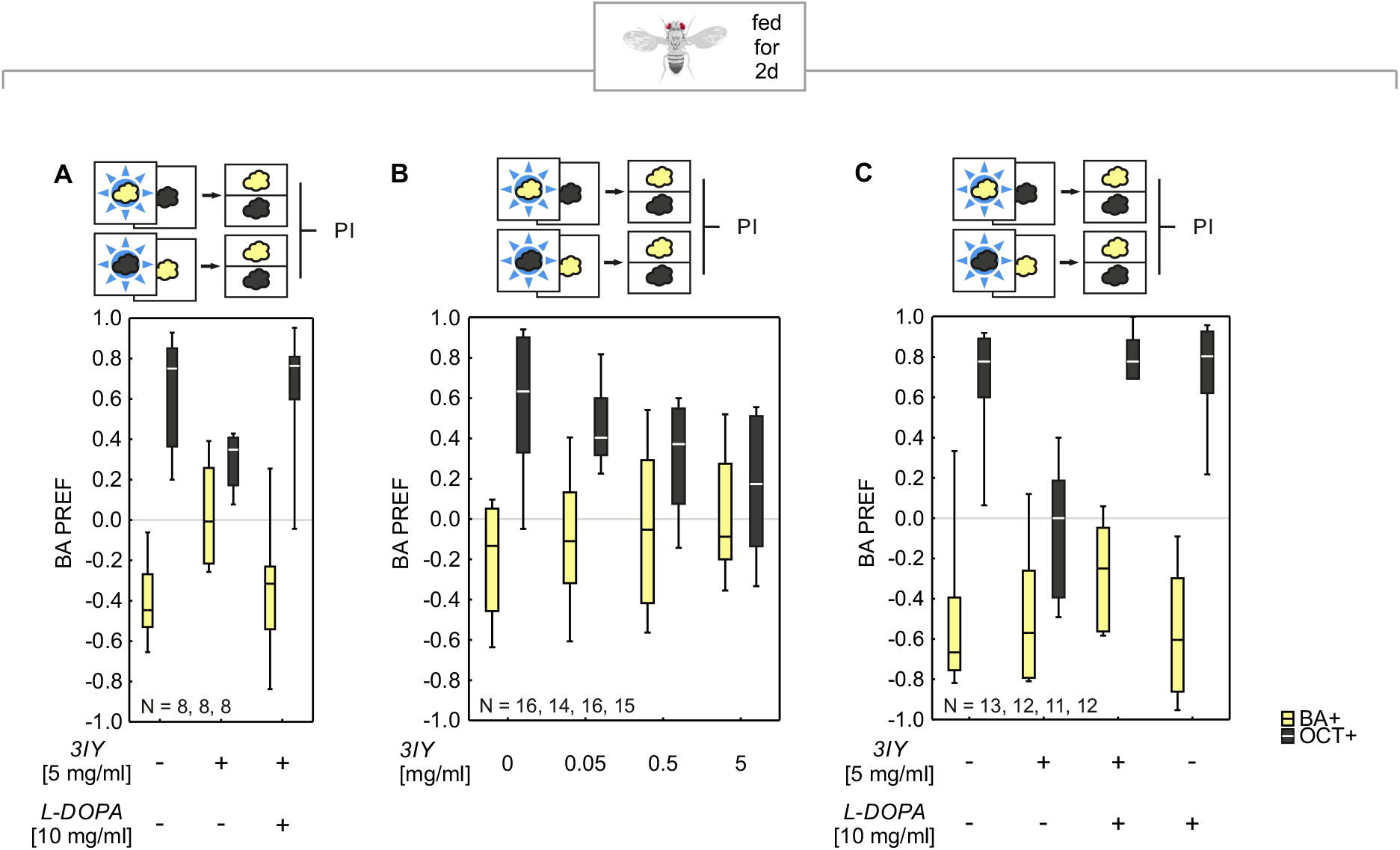
Preferences underlying the Performance Indices in Figure 4. **(A, B, C)** Preference data refer to the Performance Indices shown in Figure 4A, B and E, respectively. Yellow boxes represent BA preference after BA-paired training (BA+), black boxes after OCT-paired training (OCT+). Box plots represent the median as the midline, 25 and 75% as the box boundaries, and 10 and 90% as the whiskers. Sample sizes are indicated within the figure.

